# Hippocampal ripples during offline periods predict human motor sequence learning

**DOI:** 10.1101/2024.10.06.614680

**Authors:** Pin-Chun Chen, Jenny Stritzelberger, Katrin Walther, Hajo Hamer, Bernhard P. Staresina

## Abstract

High-frequency bursts in the hippocampus, known as ripples (80-120 Hz in humans), have been shown to support episodic memory processes. However, converging recent evidence in rodent models as well as human neuroimaging suggests that the hippocampus may be involved in a wider range of memory domains, including motor sequence learning (MSL). Nevertheless, no direct link between hippocampal ripples and MSL has yet been established. Here, we recorded intracranial electroencephalography from the hippocampus in 20 epilepsy patients during a MSL task in which participants showed steady improvement across nine 30-second training blocks interspersed with 30-second rest (‘offline’) periods. We first demonstrate that ripple rates strongly increase during rest periods relative to training blocks. Importantly, ripple rates during rest periods tracked learning behaviour, both across blocks and across participants. These results suggest that hippocampal ripples during offline periods play a functional role in motor sequence learning and that the hippocampus may be involved in offline learning beyond episodic memory.

## Introduction

How does the human hippocampus contribute to learning and memory? Following the report of patient H.M.^1^, the distinction between hippocampus-dependent ‘declarative’ and non-hippocampus-dependent ‘non-declarative’ memory has been widely accepted^2^. However, mounting evidence has begun to challenge the exclusive role of the hippocampus in canonical forms of declarative memory (e.g., episodic memory). For instance, the hippocampus has been implicated in short-term memory^3^, working memory^4^, non-conscious forms of learning^5^ as well as motor skill acquisition^6^. Irrespective of the type of learning, another key question is at which stage of learning hippocampal contributions unfold. That is, learning can be roughly divided into online and offline components, with the former denoting on-task acquisition and retrieval, and the latter including post-acquisition rest periods, spent either awake or asleep. Interestingly, accumulating evidence points to an important role of hippocampal engagement during offline periods, even in tasks that seemingly do not require the hippocampus during acquisition^7-9^.

Hippocampal neurons fire in large synchronous population bursts called sharp waves superimposed by high-frequency bursts named ‘ripples’. Initially observed in rodent hippocampus, ripples during offline states have been linked to replay of navigational trajectories along with consolidation of spatial memories^10^. In humans, hippocampal ripples can be reliably observed via direct recordings from patients undergoing invasive monitoring for the surgical treatment of refractory epilepsy^11^. In episodic memory tasks, human ripples have been associated with memory performance both during online recall^12,13^ and offline sleep periods^14^. Beyond episodic memory tasks though, the involvement of hippocampal ripples in learning remains unclear.

In the present study, we ask whether hippocampal ripples play a functional role in motor skill learning. To this end, we examined hippocampal ripple attributes during a motor sequence learning paradigm in epilepsy patients. We report a strong increase of ripple rates during offline rest relative to online task periods. Importantly, these offline ripple increases were directly linked to learning behaviour, tracking performance improvements within and across participants.

## Materials and methods

### Participants

Intracranial electroencephalography (iEEG) recordings from the human hippocampus were obtained from 20 participants (11 male, mean age 30.5 years, range 20 – 58 years, 2 left-handed; see Supplementary Table 1) undergoing invasive monitoring as part of their treatment for refractory epilepsy at the University Hospital Erlangen (Erlangen, Germany). Ethical approval was granted by the ethics commission of the Friedrich-Alexander Universität Erlangen-Nürnberg (142_12B). Written informed consent was obtained in accordance with the Declaration of Helsinki.

### Behavioural task

Participants performed an explicit motor sequence learning (MSL; Figure 1a) task^15^ with their non-dominant hand where they repetitively typed a 5-item numerical sequence (1-4-2-3-1) displayed on a screen as quickly and as accurately as possible. Keypress 1 was performed with the little finger, keypress 2 with the ring finger, keypress 3 with the middle finger, and keypress 4 with the index finger. Individual keypress times and identities were recorded for behavioural data analysis. Participants performed the typing (‘training’) task for 9 consecutive blocks, with each block lasting 30 seconds. 30 seconds rest periods were interleaved between training blocks (8 blocks of rest periods). During the training task, participants were instructed to fixate on the five-item sequence continuously displayed on the screen. To facilitate sequential training, a hexagon was shown below the to-be-pressed item and would only proceed to the next item upon a correct key press. During the rest periods, the sequence was replaced with a countdown proceeding from 30 to 0 in one-second increments, and participants were instructed to fixate on this countdown. The experiment was designed and delivered using MATLAB (MathWorks) and Psychophysics Toolbox v.3^16^. Before the task started, a practice session was administered using the simplified sequence 4-3-2-1-4. For an exploratory analysis, 8 participants were randomly assigned to a “sound” condition, whereas 12 were assigned to the “no sound” condition. In the “sound” condition, keypress 1-2-3-4 were paired with the tones CDEF, respectively. A linear mixed effects analysis revealed no effects of sound condition on learning rates (F_(1,158)_ = 1.104; *p* = 0.295). Similarly, a linear mixed effects analysis revealed no effects of handedness on learning rates (F_(1,158)_ = 0.509; *p* = 0.295). Therefore, participants were pooled together for all subsequent analyses.

**Figure 1.**
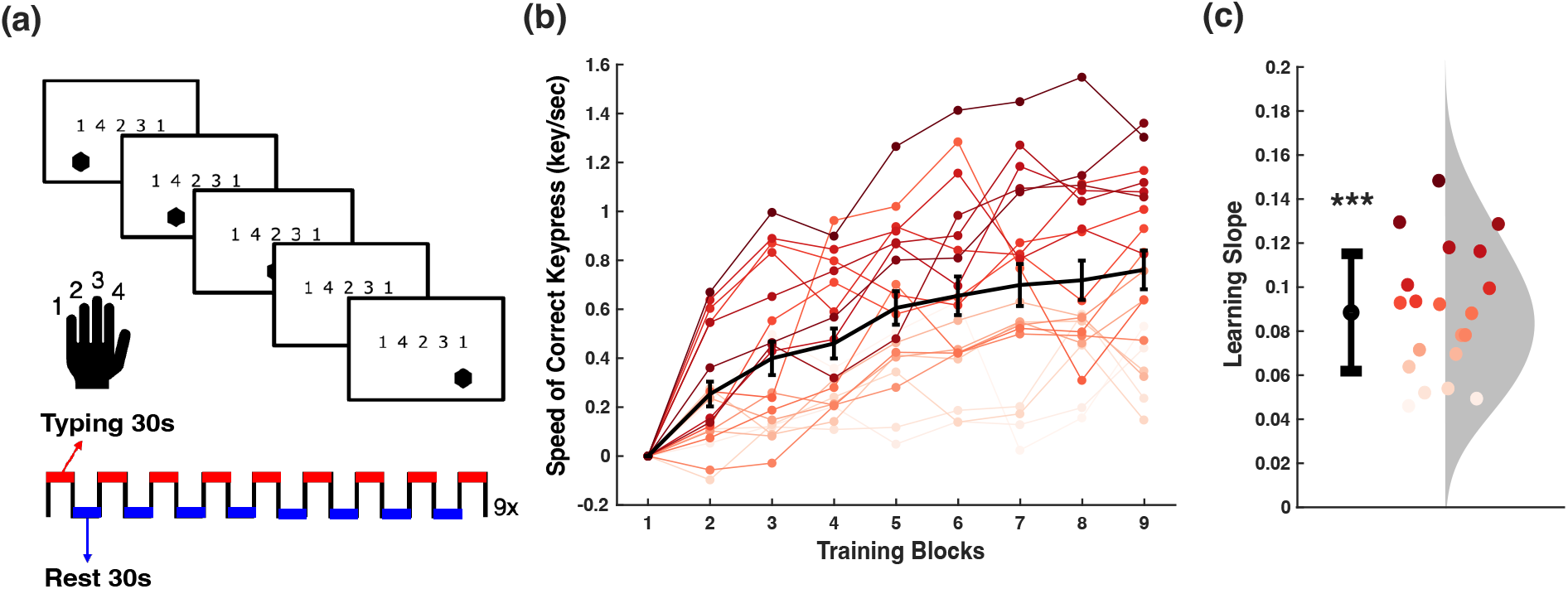
Behavioural task and performance. **(A)** Motor sequence learning (MSL) task. Participants performed an explicit motor sequence learning (MSL) task with their non-dominant hand where they repetitively typed a 5-item numerical sequence (1-4-2-3-1) displayed on a screen as quickly and as accurately as possible. Participants performed the typing task (‘training’) for 9 blocks, with each block lasting 30 seconds. 30 seconds rest periods were interleaved between training blocks (8 blocks of rest periods). **(B)** Typing speed across training blocks. Block performance was summarized as the median typing speed over the 30-seconds training block. Each line represents one participant’s performance across 9 blocks. Error bars represent standard error of the mean. For visualisation, performance of block 1 was subtracted from the remaining blocks. **(C)** Learning slope per participant. Individual slope per participant derived from the linear-mixed effects modelling (***… *p* < 0.001). Each point represents one participant’s learning slope, i.e., the magnitude of speed improvement per one training block increase. Error bars represent standard error of the mean.

### Learning metric

Motor sequence performance was quantified integrating speed and accuracy, using the inverse of time (in seconds) required to complete a correct keypress. A keypress was deemed correct if it occurred within any correct 5-item response pattern (1-4-2-3-1, 4-2-3-1-1, 2-3-1-1-4, 3-1-1-4-2, 1-1-4-2-3). This approach allowed us to quantify motor sequence learning within each training block at the resolution of individual keypresses. Block performance was summarized as the median typing speed over the 30-seconds training block^17^.

### Recording system and electrodes

Recordings were performed using Behnke–Fried depth electrodes (Ad-Tech Medical Instrument Corporation). The electrodes were connected to an ATLAS system (Neuralynx) and recorded with a sampling rate of 2048 Hz. Electrode placements were determined by clinical criteria and only patients with hippocampal electrodes were included in the current study. We used a bipolar referencing scheme using the two most medial electrode contracts. For visualization, electrode coordinates were converted to MNI space (see Figure 1) using the brainstorm-toolbox^18^.

### Ripple detection

After pre-processing steps (see Supplementary Methods), ripples were identified by an initial time domain detection procedure followed by a frequency domain assessment for accepting/rejecting candidate events. Analytic criteria were based on prior human hippocampal ripple studies^19-21^, as detailed in Supplementary Methods. For identified ripples, we summarized four attributes: 1) ripple rate (Hz), calculated by using the number of detected ripple events divided by artifact-free experiment time in seconds, 2) ripple duration (ms), 3) ripple max amplitude (mV) calculated using the ripple band-passed signal, and 4) ripple peak frequency (Hz).

### Statistical analysis

To capture the improvement of typing speed across learning blocks, linear mixed-effects models were employed, using participant and block number as random effects [*Typing Speed*_*ij*_ *∼ Block*_*j*_ *+ (Block*_*ij*_ | *Participant*_*i*_)]. Likewise, for ripple attribute-based analyses, linear mixed-effects models (using participant as a random effect) were employed to account for the nesting of multiple contacts within each participant. To correlate rest ripple rates with behaviour, resting ripple rates were first normalised as follows: *(resting ripple rates – training ripple rates) / (resting ripple rates + training ripple rates)*.

Next, a linear mixed-effects model was built to examine the relationship between ripple rates and task performance (typing speed in the subsequent block) at the block level, with normalised rest ripple rates as a fixed effect, participant as random effects, and training speed in the previous block as a covariate, accounting for individual differences in previous block performance (taking into account that better performance naturally leaves less room for improvement).

Statistical analyses were carried out using R statistical software^22^. Restricted Maximum Likelihood (REML) estimations were used to estimate variance components of the linear mixed effects models, as they can reliably produce unbiased estimators^23^. Likelihood Ratio (LR) Tests were used to compare the model of interests against its reduced model to test the significance of the model fit.

## Results

### Typing speed improvements across training blocks

To test whether performance improved across training blocks, we used a group linear mixed-effects model of typing speed, with block number as fixed effects, and block number and participant as random effects. This allowed us the examine the learning slope at the group level (fixed effects) as well as at the individual participant level (random effects). Results confirmed that participants’ typing speed reliably improved across blocks, with an increase of 0.09 correct key presses per block (F_(1,159)_ = 89.58; *p* < 0.001) (Figure 1b). Random effects confirmed that participants’ learning slopes were positive (Figure 1c), showing that all participants’ typing speed improved across training blocks.

### Ripple attributes for MSL training and rest periods

Hippocampal ripples were detected based on established criteria after rigorous artifact rejection (see Supplementary Methods). Figure 2b shows the grand-average ripple across *n* =38 contacts along with an individual ripple example. After ripple events were detected, we examined four attributes of ripples during MSL training vs. interspersed resting periods: rate, duration, amplitude, and peak frequency (with Bonferroni correction for multiple comparisons). Results revealed an increase of ripple rate during resting compared to training periods. Figure 2c illustrates this effect by showing a raster plot of ripples pooled across all contacts, participants and training vs. resting periods. Group linear mixed-effects analysis of ripple attributes, with task condition (training, resting) as a fixed effect and participant as random effects, revealed a significant main effect of task for ripple rate, where ripple rates were higher during resting periods than during training periods by 0.01 Hz on average (Figure 2d, F_(1,50)_ = 24.250; *p* < 0.001). Ripple duration was longer during resting compared to training periods by 1.336 ms (F_(1,50)_ = 4.098; *p* = 0.048), but this effect did not survive Bonferroni correction. No statistical differences were seen in amplitude (F_(1,50)_ = 0.012; *p* = 0.914) and frequency (F_(1,50)_ = 0.340; *p* = 0.855).

**Figure 2.**
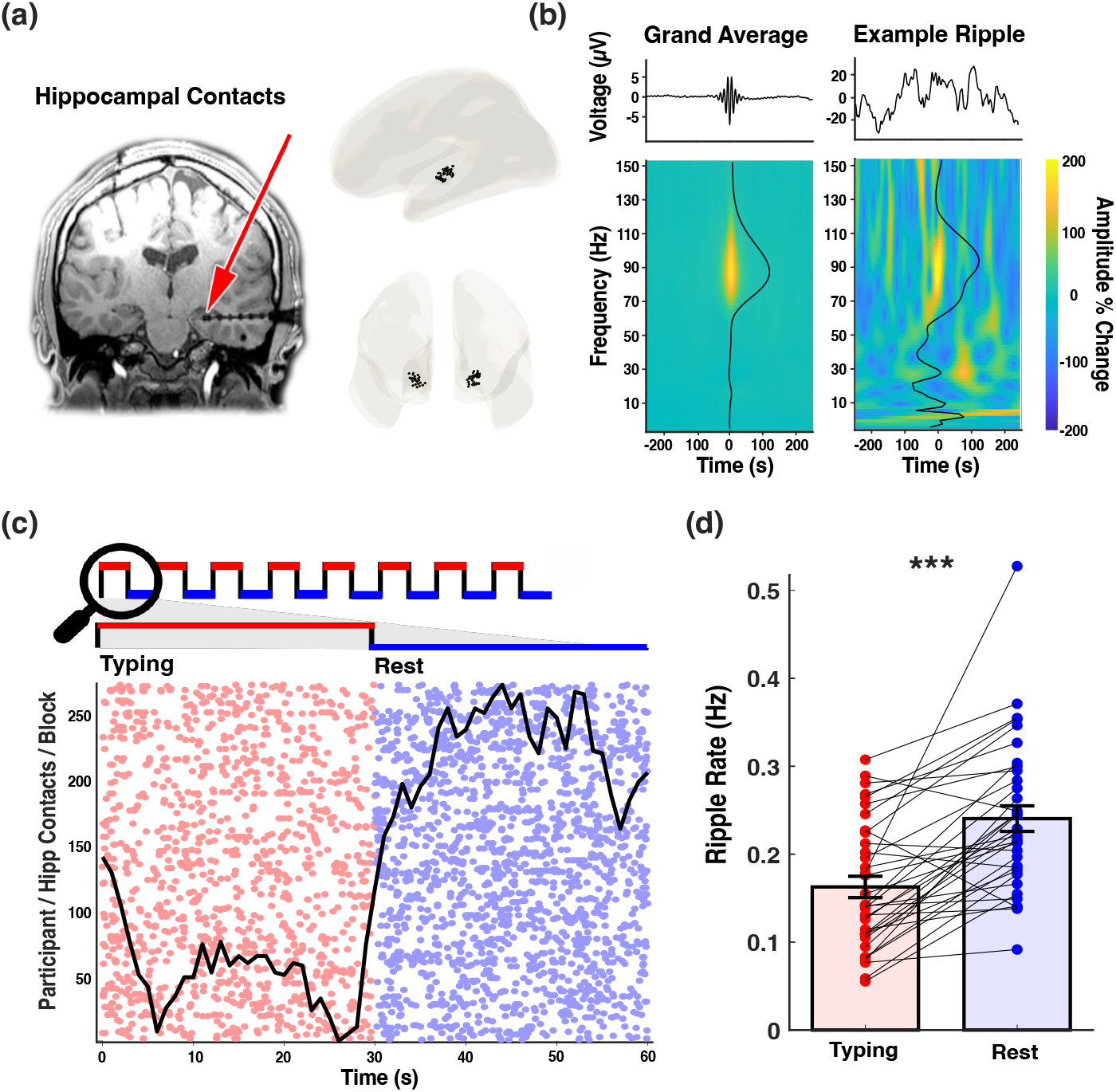
Hippocampal ripples modulated by task condition. **(A)** Hippocampal contacts. Left panel: anatomical location of an example electrode with two contacts in the hippocampus. Right panel: Hippocampal contacts from all participants rendered on a brain template in MNI space. **(B)** Hippocampal ripples. *Left*: grand average of ripple-locked raw voltage trace and time-frequency decomposition from all ripples detected during the experiment across all participants. *Right*: ripple-locked raw voltage trace and time-frequency decomposition of an example ripple. Colour of the time-frequency plot represents amplitude change compared to pre-event baseline (−1.5 to -0.5 from ripple centre). Black line on the time-frequency plot represents amplitude density across frequency bands. **(C)** Raster plots of ripple occurrences during MSL typing and rest periods, illustrating the ripple rate increase during rest periods. Each row represents one block of training and rest periods from each contact/participant. **(D)** Bar graph of summarized ripple rate during MSL typing and rest periods. Ripple rates were higher during rest than during typing periods. Each point represents the average ripple rate for each bipolar referenced contact (***… *p* < 0.001).

### Link between offline ripples and learning behaviour

Is the increase in hippocampal ripples during offline periods directly linked to motor sequence learning? If so, ripple rates during resting periods should mirror behavioural performance increases across blocks (Figure 1b). Examining training and resting ripple rates across blocks, we indeed observed a steady increase in resting ripple rates (Figure 3a). To quantify this effect, we used a group linear mixed-effects analysis to examine the interaction between MSL experiment blocks (1-8) and task condition (training vs rest), using task condition and block number as fixed effects and participant as random effects. Results revealed a significant interaction between task condition and block number on ripple rate (F_(1,524)_ = 6.109; *p* = 0.014), due to the difference between resting vs. training ripple rates increasing by 0.01 Hz on average per block. We tested the model against its nested model (only including main effects of condition and block, without the interaction term), using a Likelihood Ratio (LR) Test, which confirmed that the interaction significantly contributed to the modulation of ripple rate (LR = 6.108; *p* = 0.014). To more directly test the association between offline ripple rates and performance increases across blocks, we modelled the slope of typing speed and the slope of normalised rest ripple rates across learning blocks using linear mixed-effects models, with blocks as a fixed effect and block/participant as random effects. We then extracted the slope term for each participant and correlated the normalised resting ripple slopes with typing speed slopes. Across participants, we observed a positive correlation (Figure 3b; Spearman’s rho = .556; *p* = .022). In short, participants who showed a greater performance increase across blocks of training also had higher ripple rate increases across blocks of resting periods.

**Figure 3.**
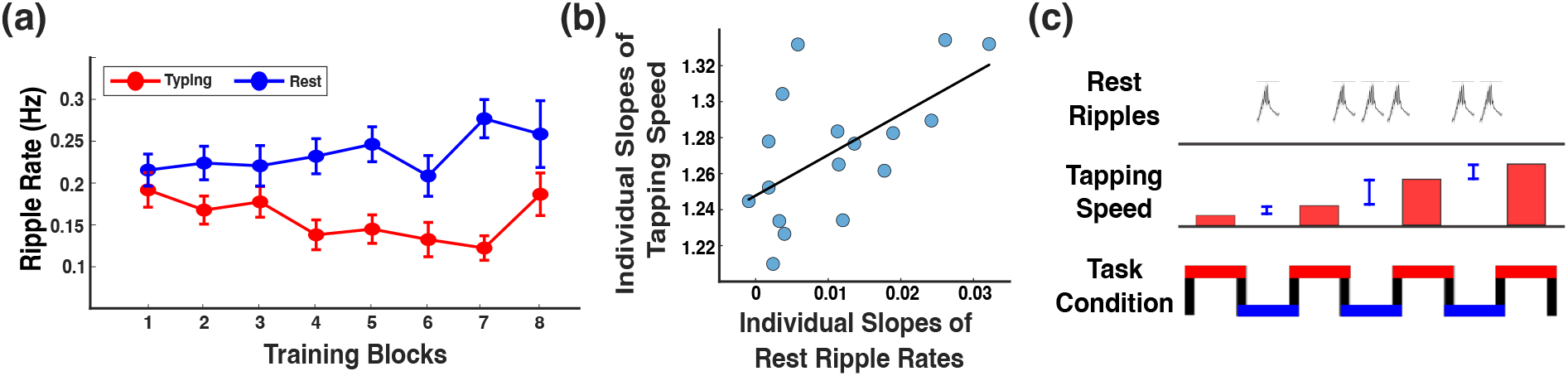
Link between ripples and learning behaviour. **(A)** Ripple rate as a function of experiment blocks and task condition (typing vs. rest). As the block number increases, the ripple rate difference between rest vs. typing periods increases. **(B)** Individual learning slopes (typing speed across blocks) correlate with rest ripple rate slopes across participants (Spearman’s rho = .556; *p* = .022). **(C)** Schematic depiction of the finding that rest ripple rate predicts block-to-block improvements in typing speed. *Bottom*: Experimental task structure, with training blocks (red) interleaved with resting periods (blue). *Middle*: Typing speed symbolised by red bars and block-to-block improvement in typing speed symbolised by blue vertical lines. *Top*: ripples rates during resting blocks track block-to-block improvements in typing speed.

Lastly, we wanted to test whether offline ripples not only track MSL performance across participants, but also predict – within a given participant – block-to-block changes in performance (Figure 3c). In other words, are stronger performance improvements from block *n* to block *n*+1 accompanied by greater ripple rates in the intermittent resting period? To tackle this question, we used a group linear mixed-effects analysis to examine whether the normalised ripple rates during rest periods (ripple rate during resting compared to training) can predict typing speed, with block number and participant as random effects. We used typing speed in the previous block as a covariate, allowing to isolate whether changes in typing speed between blocks are predicted by ripple rates during interleaved rest periods. Group linear mixed-effects analysis revealed a significant positive relationship between offline ripple rate and typing speed improvement, such that every unit increase in normalised ripple rates is associated with a 0.08 keypresses/second increase in typing speed (F_(1,231)_ = 12.048; *p* < 0.001). We tested the model against its nested model (only including typing speed in the previous block as the fixed effects; without ripple rate), using a LR Test, and found that normalised rest ripple rates significantly contributed to the prediction of typing speed during the subsequent block (LR ratio = 5.64; *p* = .018).

## Discussion

Hippocampal ripples have recently emerged as a viable mechanism to support episodic memory processes in humans. However, beyond episodic memory, we have the remarkable ability to continuously acquire complex motor skills, from riding a bicycle to playing a musical instrument. Before becoming effortless and automatic, this kind of learning involves integrating separate movements into a unified and coordinated sequence of actions under certain rules or constraints. Here, using a well-established motor sequence learning (MSL) task (Figure 1), we asked whether the hippocampus might be involved in learning such motor skills. We first confirmed that all participants improved performance across experimental blocks. Next, we demonstrated that hippocampal ripple rates were strongly modulated by MSL resting vs. typing periods (Figure 2). Finally, we found direct links between these offline ripple rates and learning (Figure 3). Specifically, rest period (vs. typing) ripple rates not only increased with block number, but also predicted performance changes across blocks as well as across participants. These findings suggest that hippocampal ripples during rest periods contribute to motor sequence learning.

A recent human iEEG study examined hippocampal ripples during various cognitive tasks and found that ripples were not unique to episodic memory behaviour^21^. Instead, ripples occurred with similar attributes during other visual perceptual tasks. Of note, ripples showed highest occurrence rates and longest durations during resting states (6-min eyes open/closed) compared to active tasks. These results dovetail with evidence that the hippocampus contributes to learning during offline periods even in tasks that show no apparent hippocampal involvement during active acquisition^8,9^. It deserves mention though that recent work in humans has also implicated hippocampal ripples during ‘online’ parts (encoding and retrieval) of episodic memory tasks^12,13^, albeit without assessing ripples during offline delay periods in the same experiment. Our current results suggest that at least in motor skill acquisition, offline ripples might play a more important role for behavioural improvements than online ripples.

How might hippocampal ripples during offline periods contribute to motor skill learning? A recent model proposes that the hippocampus may act as a sequence generator, connecting encoded items of different modalities in space/time via ripples^10^. We speculate that during MSL rest periods, the hippocampus may reactivate information about the sequence, leading to behavioural improvements in task performance. The relevance of offline periods for motor skills acquisition has been well established. For example, distributed practice, where frequent rest periods are interspersed with practice (training) blocks, has been shown to be more effective for skill learning compared to massed practice, where the same total amount of practice is performed over longer continuous blocks^24^. In light of our current findings, one tentative interpretation of this pattern is that an increased number of offline periods provides more opportunity for the hippocampus to drive reactivation and set consolidation processes in motion. It is interesting to note that ripples increased not only in occurrence during offline periods in our paradigm, but also in duration (although the effect did not survive Bonferroni correction). Ripple duration has previously been associated with memory reactivation^25^ and hippocampal-cortical communication during sleep^20^. Indeed, human fMRI work has provided evidence of hippocampal reactivation after motor skill learning and its relation to consolidation^26^. Moreover, recent studies employing fine-grained block-by-block analyses of MSL tasks (‘micro-online and -offline gains’) have shown that interspersed rest periods promote consolidation at much shorter timescales (i.e., seconds and minutes) than previously thought^27-29^. Accordingly, the ripple engagement we observed during MSL rest periods may reflect a form of rapid consolidation via reactivation that contributes to behavioural improvements across training.

While our work focused on hippocampal ripples, motor sequence learning most likely requires dynamic interactions between the hippocampus with cortical (e.g., motor cortex) as well as other subcortical regions (e.g., striatum). Indeed, using multivariate decoding of magnetoencephalography (MEG) data, spontaneous replay of the motor sequence representation was reported in sensorimotor and hippocampal regions during wake rest periods, which predicted the magnitude of rapid consolidation^17^. Future work combining hippocampal recordings with systematic coverage of specific cortical areas will be needed to understand how hippocampal ripples may drive the cortical replay of motor sequences.

In conclusion, our findings integrate and extend prior work in the human and animal hippocampus to suggest that hippocampal ripples during offline periods are associated with motor skill learning. This finding adds growing support to the view that hippocampal ripples serve as an internally generated and state-dependent mechanism for learning beyond the episodic memory domain.

## Data availability

Derivative data, MATLAB and R scripts, and results presented in all figures will be publicly available on the Open Science Framework (https://osf.io/9r8pu/) upon publication.

## Supplementary material

### Supplemental Methods

#### Artifact rejection

An automated algorithm was applied to identify artifacts for each bipolar channel based on three versions of the raw data. In each case, artifactual samples were defined as exceeding the median + 4 * interquartile range (IQR) across all time points: (i) absolute amplitudes of the raw signal, (ii) absolute amplitude of the first derivative of the raw signal (i.e., gradient artifacts likely caused by interictal spikes) and (iii) amplitude of the root mean square (RMS; 100 ms window) after high-pass filtering the signal at 250 Hz. Next, we employed two additional measures to exclude epileptogenic activity. First, based on the sum power across the frequencies 1–60 Hz (30 logarithmically spaced frequencies; time-frequency decomposition using Morlet wavelets with seven cycles, followed by taking the natural logarithm and frequency-specific z-scoring across time) exceeding 4 IQR above the median sum power. The rationale behind these criteria was that epileptogenic activity exhibits high amplitudes, sharp amplitude changes, and power increases across a broad frequency range. Lastly, we employed an automatic interictal spike detection algorithm^1^. All detected artifact samples were then padded by ±1 second (see Supplementary Fig. 1). Furthermore, artifact-free intervals shorter than 1s were also marked as artifacts. The thresholds used are based on previous publications^2-5^ and automated detection was followed by visual inspection. Contacts with more than 2/3 of task duration (out of 9 blocks of 30s training + 8 blocks of 30s rest) contaminated with padded artifacts were excluded. Consequently, four bipolar-referenced contacts from three participants were excluded from ripple analyses (final contacts *n* = 34; participants *n* = 17).

**Supplemental Figure 1.**
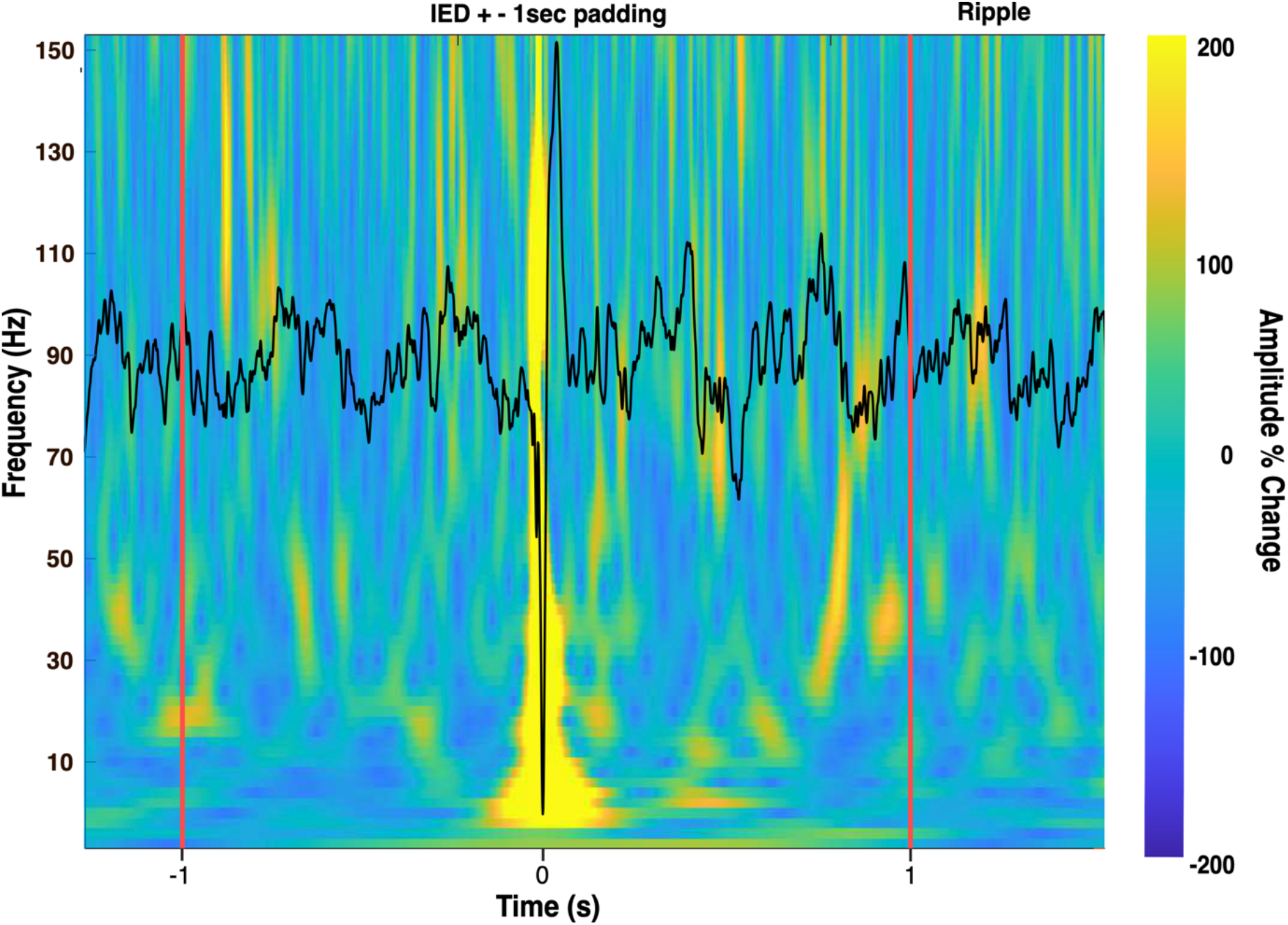
Hippocampal ripples modulated by task condition. Illustration of a detected IED and 1-second padding. Ripple candidates detected within the padded window were rejected. Only the ripple after 1-second padding was considered in this figure.

#### Ripple detection and rejection

Signals from hippocampal contacts (i.e., continuous bipolar re-referenced time-series) were first band-pass filtered from 80 to 120 Hz (ripple band) using a 4^th^-order FIR filter. Next, the RMS of the band-passed signal was calculated and smoothed using a 20-ms window. Ripples were detected based on amplitude and duration thresholds of this RMS time course. Specifically, ripple events were identified as having an RMS amplitude above 1.5, but no greater than 9 standard deviations from the mean. Ripple duration was defined as the supra-threshold time of the RMS signal. Detected ripple events with a duration shorter than 38 ms (corresponding to 3 cycles at 80 Hz) or longer than 500 ms were rejected. In addition to the amplitude and duration criteria the spectral features of each detected ripple event were then examined^4,5^: 1) candidate ripple events were rejected if the most prominent peak was outside the ripple band (80-120Hz); 2) candidate ripple events where peaks within high frequency activity (30-200Hz) that are outside of ripple band (80-120Hz) exceed 80% of the ripple peak height; 3) candidate ripple events with more than 1 peaks in the upper-frequency range (120-200 Hz) were rejected; 4) candidate ripple events were rejected if ripple-peak width was greater than 3 standard deviations from the mean ripple-peak width calculated for a given electrode. On average, 23.4% (SD = 9.9%) of candidate ripples during the experiment were rejected.

### Supplemental Table

**Table 1.** Participant demographic and electrode information. Demographic and electrode information is reported for each subject (1-20), including sex (male/female), age at time of experiment (years), total number of hippocampal contacts.

| Participant | Sex | Age | Handedness | # bipolar ref contacts |  |  |
| --- | --- | --- | --- | --- | --- | --- |
|  |  |  |  | Tot. | L | R |
| 1 | M | 21 | R | 2 | 2 | 0 |
| 2 | M | 20 | R | 0 | 0 | 0 |
| 3 | M | 30 | R | 2 | 0 | 2 |
| 4 | F | 20 | R | 2 | 0 | 2 |
| 5 | F | 47 | R | 2 | 2 | 0 |
| 6 | M | 39 | R | 2 | 0 | 2 |
| 7 | F | 58 | L | 2 | 1 | 1 |
| 8 | F | 23 | R | 2 | 2 | 0 |
| 9 | M | 39 | R | 2 | 0 | 2 |
| 10 | F | 36 | R | 2 | 2 | 0 |
| 11 | M | 19 | L | 2 | 0 | 2 |
| 12 | M | 23 | R | 0 | 0 | 0 |
| 13 | F | 35 | R | 2 | 0 | 2 |
| 14 | F | 24 | R | 2 | 2 | 0 |
| 15 | F | 35 | R | 2 | 2 | 0 |
| 16 | M | 26 | R | 2 | 2 | 0 |
| 17 | F | 38 | R | 0 | 0 | 0 |
| 18 | F | 29 | R | 2 | 2 | 0 |
| 19 | M | 27 | R | 2 | 2 | 0 |
| 20 | M | 21 | R | 2 | 0 | 2 |

